# Dynein catch bond as a mediator of codependent bidirectional cellular transport

**DOI:** 10.1101/250175

**Authors:** Palka Puri, Nisha Gupta, Sameep Chandel, Supriyo Naskar, Anil Nair, Abhishek Chaudhuri, Mithun K. Mitra, Sudipto Muhuri

**Author notes:** Electronic mail (Abhishek Chaudhuri). Electronic mail (Mithun K. Mitra). Electronic mail (Sudipto Muhuri).

## Abstract

Intracellular bidirectional transport of cargo on Microtubule filaments is achieved by the collective action of oppositely directed dynein and kinesin motors. Experiments have found that in certain cases, inhibiting the activity of one type of motor results in an overall decline in the motility of the cellular cargo in both directions. This counter-intuitive observation, referred to as paradox of codependence is inconsistent with the existing paradigm of a mechanistic tug-of-war between oppositely directed motors. Unlike kinesin, dynein motors exhibit catchbonding, wherein the unbinding rates of these motors decrease with increasing force on them. Incorporating this catchbonding behavior of dynein in a theoretical model, we show that the functional divergence of the two motors species manifests itself as an internal regulatory mechanism and provides a plausible resolution of the paradox of codependence.

---

Bidirectional transport is ubiquitous in nature in the context of intracellular transport^1–4^. Within the cell, oppositely directed motor proteins such as *dynein* and *kinesin* motors walking on microtubule (MT) filaments^1,5^ transport diverse organelles like mitochondria, phagosomes, endosomes, lipid droplets and vesicles. While the phenomenon of bidirectional transport has been well studied experimentally under both *in-vitro* and *in-vivo* conditions for variety of different systems^1,6^, the underlying mechanism by which the motors involved in bidirectional transport are able to achieve regulated long distance transport is far from clear and is a subject of much debate^1,2,6–10^.

A theoretical framework proposed to explain the bidirectional transport is based on the *tug-of war* hypothesis^1,3,5,6,9,11,12^. The basic underlying premise of this hypothesis is that the motors act independently, stochastically binding to and unbinding from the filament and mechanically interacting with each other through the cargo that they carry (Fig. 1a)^6,9,11^. The resultant motion arises due to the competition between the oppositely directed motors with the direction of transport being determined by the stronger set of motors^9,11^. While many experiments have provided support for this mechanical tug-of-war picture^5,9,13–15^, there remain a large class of experiments whose findings are incompatible with the predictions of this model^6,16–21^

The *tug-of-war* model predicts that the mechanical competition between the two motors would lead to an enhancement of motility in one direction on inhibiting the activity of the other motor. However, a range of experiments have shown that there exists some coordination mechanism due to which inhibition of one motor species results in an overall decline in the motility of the cargo^2,6,17–19^. This apparently counterintuitive finding has been referred to as the *paradox* of *codependence*, suggesting some kind of coordination between the oppositely directed motors which has not been accounted for in the theoretical *tug-of-war* model^1,6^. An open question is then how this paradox can be resolved and understood in terms of the underlying mechanism which governs bidirectional transport. In this work we seek to address this issue by re-examining the theoretical *tug-of-war* model^9,11^.

A striking difference between the single molecular behaviour of dynein and kinesin lies in their unbinding kinetics. Unlike kinesin, dynein can exhibit catchbonding, where the propensity for the dynein motors to unbind from cellular filament decreases when subjected to increased load force (Fig1b)^10,22,23^. In contrast, the detachment rate of kinesin motors increases exponentially with increasing load force - a characteristic of *slip bond*^8,23,24^. The effect of dynein catchbonding on bidirectional transport has been investigated theoretically in context of studying bidirectional transport of lipid droplets under both *in-vitro* and *in-vivo* conditions^10^. A detailed quantitative comparison of theoretical predictions of typical cargo trajectories, and other transport characteristics such as pause durations, time between pauses, runlengths with experimentally obtained data has shown significant divergence. Different mechanisms such as the effect of regulatory proteins like JIP1^25^ or the finding that the cargoes might have some memory^23^ have been proposed to explain the quantitative discrepancy between experimental observations and theoretical predictions. While these studies demonstrate that catch-bonding mechanism *alone* cannot explain the quantitative features of bidirectional transport characteristics of lipid droplets, delineating the role of catchbonding itself as a mediator of codependent bidirectional transport has not been investigated and probed.

**FIG. 1.**
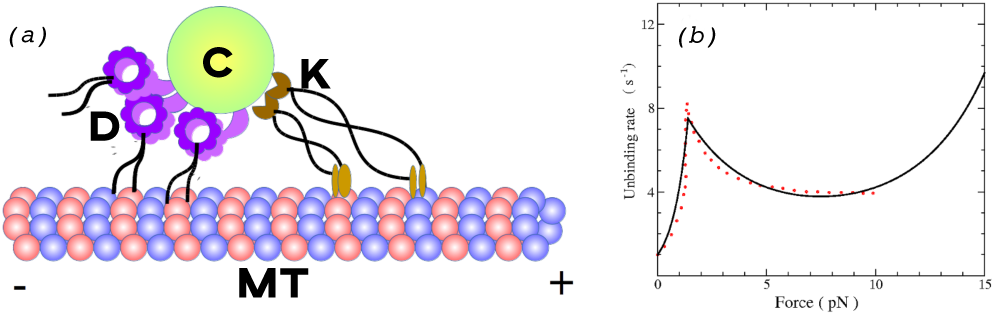
(a) Schematic of bidirectional motion of cargo (C) attached to both kinesin (K) and dynein (D) motors on a microtubule (MT) filament; (b) Single dynein unbinding rate from experiments^10^ (points) and the corresponding fit (solid line) from the TFBD model^26^.

In this paper, we specifically focus our attention on the emergence of codependent transport characteristics due to the effect of the dynein catch bond. We use a threshold force bond deformation (TFBD) model, which correctly reproduces collective transport properties of unidirectional transport^26^, to fit the experimentally observed unbinding rate of single dynein motors. With the TFBD model for dynein, and the usual slip bond model for kinesin^8,9^, we study the transport properties of bidirectional cargo motion by multiple motors. We use experimentally relevant measures to characterize the transport properties of cellular cargo: (i) average processivity, defined as the mean distance a cargo travels along a filament before detaching, and (ii) probability distributions of runtimes and pause times. We show that in biologically relevant parameter regimes, catchbonding significantly alters the transport characteristics. Further, our study shows the existence of an internal regulatory mechanism of transport due to dynein catch bonding and its importance in fashioning the codependent transport behaviour. Indeed a complete description of cellular transport necessarily involves an internal regulatory mechanism mediated by the catch bond a one of the necessary ingredients, in addition to external regulatory factors.

## RESULTS

We study transport of a cellular cargo with *N*_+_ kinesin motors and *N*_−_ dynein motors. Each of these motors stochastically bind to a MT filament with rates π_±_ and unbind from the filament with rates *ε*_±_. The instantaneous state of the cargo is expressed in terms of the number of kinesin (0 ≤ *n*_+_ ≤ *N*_+_) and dynein (0 ≤ *n*_−_ ≤ *N*_−_) motors that are attached to the filament. At any instant only the attached set of motors generate force on the cargo and are involved in its transport.

The load force is assumed to be shared equally among the attached motors. We use the Stochastic Simulation Algorithm (SSA)^27,28^ to obtain individual cargo trajectories governed by the associated Master Equation (see Section A for details) for the probabilities in state space of attached kinesin and dynein. The cargo is considered to detach from the MT filament when *n*_+_ = *n*_−_ = 0. The simulated trajectories are then analysed to quantify the statistical properties of the system.

Detailed experimental studies have revealed that dynein motors exhibit catchbonding at forces larger than the stall force, *F_s_*_−_, defined as the load force at which the cargo stalls^10,22,23^. This catchbonding regime is characterised by a decreasing detachment rate with increasing opposing load. The unbinding rate of a single dynein is modeled by

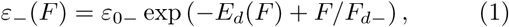

where the deformation energy *E_d_* sets in beyond the stall force, and is modeled by a phenomenological equation^26^,

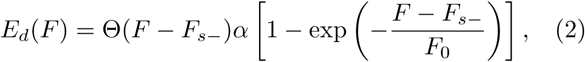

The parameter *α* sets the strength of the catch bond, while *F*_*d*−_ and *F*_0_ characterise the force scales for the dissociation energy and the deformation energy respectively. This correctly reproduces the experimentally reported dissociation dynamics of a single dynein as shown in Fig. 1(b)^26^. The unbinding kinetics of kinesin exhibits usual *slip* behavior. The characteristic stall forces and detachment forces of kinesin are denoted by *F_s_*_+_ and *F_d_*_+_ respectively. The values for the various parameter used in the stochastic simulations are listed in Table I.

The non-linear force response of the catch-bonded dynein has non-trivial implications for the transport properties of cargo during bidirectional transport. We investigate the consequences of this non-linear behaviour using the average processivity of the transported cargo and the probability distributions of cargo runtime and pausetimes in a particular direction. We relate it with the nature of individual cargo trajectories, observed in the context of various *in-vivo* and *in-vitro* experiments.

### Processivity characteristics

In Fig. 2 (a) we show the effect of variation of *N*_−_ on processivity, defined as the net displacement of the cargo until it unbinds. In the absence of catch bond (*α* = 0), for a fixed value of *N*_+_, the processivity decreases continuously with increasing *N*_−_, indicating a decreasing net movement in the positive direction, as expected from the conventional tug-of-war argument. Within the range of parameters investigated in our model, the dynein stall force has no effect on transport characteristics. When catch bond is incorporated (*α* > 0), the consequences are quite dramatic. For strong dynein (*F*_*s*−_ = 7pN), the processivity in the positive direction drops significantly even for one dynein motor, almost stalling the cargo. Increasing *N*_−_ further, eventually stalls the cargo, with no movement observed in either direction. For weak dynein (*F_s_*_−_ = 1pN), increasing *N*_−_ not only stalls the cargo, but also forces it to move in the negative direction. Weak dynein switches on its catchbond at smaller values of load force, leading to an increased propensity to latch on to the filament. This results in negative-end directed motion even for a small number of dynein motors. Strong dynein does not engage its catchbond until at relatively high values of load force.

**FIG. 2.**
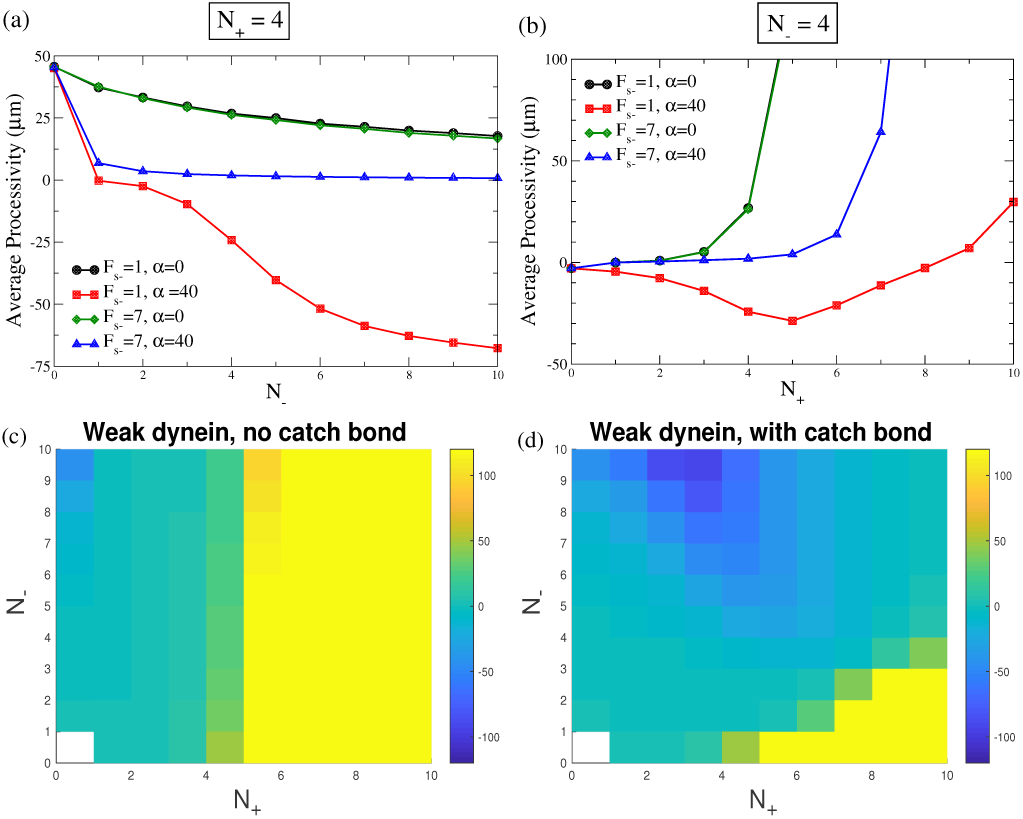
Processivity (a) as a function of *N*_-−_ for *N*_+_ = 4, and (b) as a function of *N*_+_ for *N*_−_ = 4. Contour plots for processivity in the *N*_+_ − *N*_−_ plane for (c) *F_s_*_−_ = 1pN, *α* = 0, and (d) *F_s_* _−_ = 1*pN*,*α* = 40*k_B_T*. The colorbar indicates the average processivity (in *µm*). Yellow regions denote strong plus ended runs, while dark blue regions indicate strong minus ended runs. The zero-force (un)binding rates for dynein are *ε*_0−_ = π_0−_ = 1/*s*

In Fig. 2 (b), we look at the effect of variation of *N*_+_ on processivity, for a fixed value of *N*_−_. Without catch bond (*α* = 0), *N*_+_ leads to a rapid increase of processivity in the positive direction due to the larger pull of the kinesin motors. In the presence of catch bond (*α* > 0), strong dynein behaves qualitatively in a similar fashion to that without catch bond. Weak dynein on the other hand engages its catch bond even for small load forces and is therefore pulled further in the negative direction. This leads to the striking phenomenon of increasing negative directed motion on increasing the number of kinesin motors. Beyond a certain number of kinesins, the motion in the negative direction is hindered, and eventually, for very large *N*_+_, the kinesin motors take over, leading to net positive-directed motion. This initial increase of processivity in the negative direction is a remarkable feature arising due to catchbonding in dynein, where increasing the number of motors of one type facilitates motion in the opposite direction, contrary to usual tug-of-war predictions, and is reminiscent of the *paradox of codependence*.

**FIG. 3.**
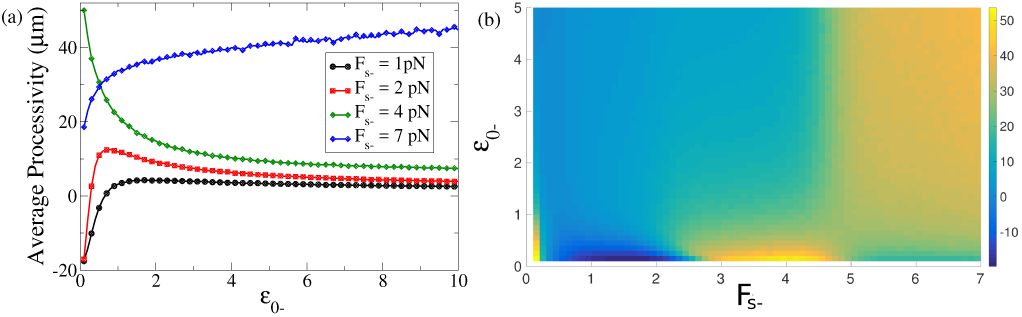
(a) Processivity as a function of *ε*_0−_ for different stall forces at *α* = 40*k_B_T*; (b) Contour plots of processivity in (*F*_*s*−_ − *ε*_0−_) plane for *α* = 40*k_B_T*. Data shown is for *N*_+_ = 6, *N*_−_ = 2, π_0−_ = 1/*s*

The corresponding contour plots of the processivity of the cargo in the (*N*_+_ − *N*_−_) plane are shown in Fig. 2(c-d), for weak dynein where the effect of dynein catch-bond is robust. As expected, in the absence of catch-bond (*α* = 0) (Fig. 2(c)), there is a smooth transition at a critical *N*_+_ from a regime where the cargo moves in the negative direction to one which moves in the positive direction. In the presence of catch-bonded dynein (Fig. 2(d)), we observe a distinct regime where the processivity increases in the negative direction on increasing *N*_+_. Plus-end directed motion occurs only in a small region of the (*N*_+_, *N*_−_) space for large *N*_+_ and low *N*_−_. The contour plot therefore provides an experimentally testable parameter space to explore the apparently anomalous codependent behavior observed in bidirectional transport when dynein catch bond is incorporated in the tug-of-war model. It is also possible to understand the effect of catch bond on processivity in terms of the average number of bound motors (see Supplementary Fig. S3 for details) which corroborate the features observed in Fig. 2.

Experimental techniques to modulate cargo processivity can also be achieved by modifying the binding/unbinding rates of the motor proteins. Dynactin mutations in *Drosophila* neurons affect the kinetics of dynein binding to the filament, leading to cargo stalls^17^. To investigate this, we tune the bare unbinding rate of dynein motor (*ε*_0−_) (Fig. 3). We observe that for weak dynein (*F_s_*_−_ = 1*pN* in Fig. 3(a)), the processivity starts to decrease from a negative value and finally saturates to a small positive value on increasing *ε*_0−_. Increasing *ε*_0−_ effectively weakens the propensity of dynein to stay attached to the filament. Beyond the critical *ε*_0−_, weakening the dynein further does not lead to any increase in the run length in the positive direction, as might be expected from a conventional *tug-of-war* scenario. At *F_s−_* = 2*pN*, on increasing *ε*_0−_, the run length in the negative direction initially decreases, then increases in the positive direction, and then decreases again, saturating to a positive value. At a larger value of the stall force (*Fs*_−_ = 4*pN*), on weakening dynein, the run length in the positive direction decreases throughout the *ε*_0−_range. This counterintuitive result is purely due to catch bonding of dynein. For strong dynein (*F_s_*_−_ = 7*pN*), we recover back the expected trend of conventional tug-of-war models, where increasing the unbinding rate consistently increases the processivity in the positive direction.

**FIG. 4.**
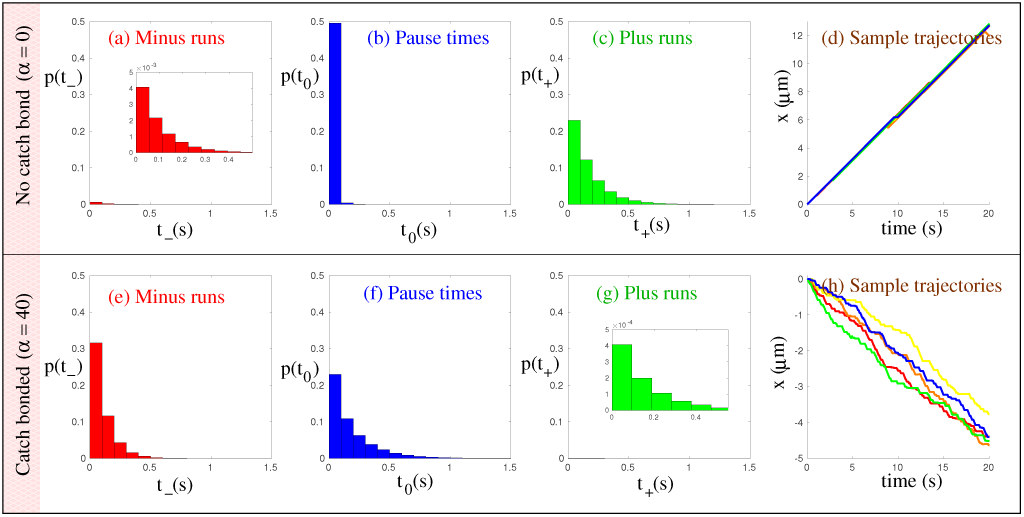
Probability distributions of runtimes for *N*_+_ = 2, *N*_−_ = 6. The top panels show the normalized histograms and sample trajectories for dynein in the absence of catch bond (*α* = 0). The bottom panels show the corresponding quantities for catch-bonded dynein (*α* = 40). (a) and (e) Distributions of runtimes for minus directed runs (shown in red); (b) and (f) pausetime distributions (shown in blue); (c) and (g) distributions of runtimes for minus directed runs (shown in green); and (d) and (h) sample trajectories. Insets, where present, show a magnified view of the probability distributions.

This entire spectrum of behaviour can be visualised as a contour plot of the processivity in the (*F_s_*_−_ − *ε*_0−_) plane (Fig. 3(b)). These contour plots capture the diversity of the processivity behaviour and codependent nature of transport due to catchbonding. For instance in Fig. 3(b) for a range of stall force for dynein *F_s_*_−_ between 1.5*pN* and 2.5*pN*, effect of increase of *ε*_0−_ can result in initial decrease in minus-end runs, leading to net positive run length. Increasing *ε*_0−_ further, leads to a reduction of the positive run length - a feature akin to reentrant behaviour^29^. The contour plot also highlights the role of the dynein stall force in determining the overall motion of the cellular cargo.

The strength of catchbond (*α*) also plays an important role in determining the nature of processivity of the cargo (see Supplementary Fig. S4). Further experiments on the exact mechanism of the catch bond in dynein can help identify biologically relevant regimes for *α* and therefore constrain the predictions of the model.

### Probability Distribution of run and pause times

Experiments on *in-vitro* and *in-vivo* systems of endosome motion have established that there is often an asymmetry in the number of motors that are simultaneously attached to a cargo. The role of catchbonding in determining the specific nature of bidirectional transport and corresponding cellular cargo trajectories can be understood by analyzing the the probability distributions of the time the cargo spends in the paused (*tug-of-war*) state versus the time it spends in the moving plus-end directed and minus-end directed state.

**FIG. 5.**
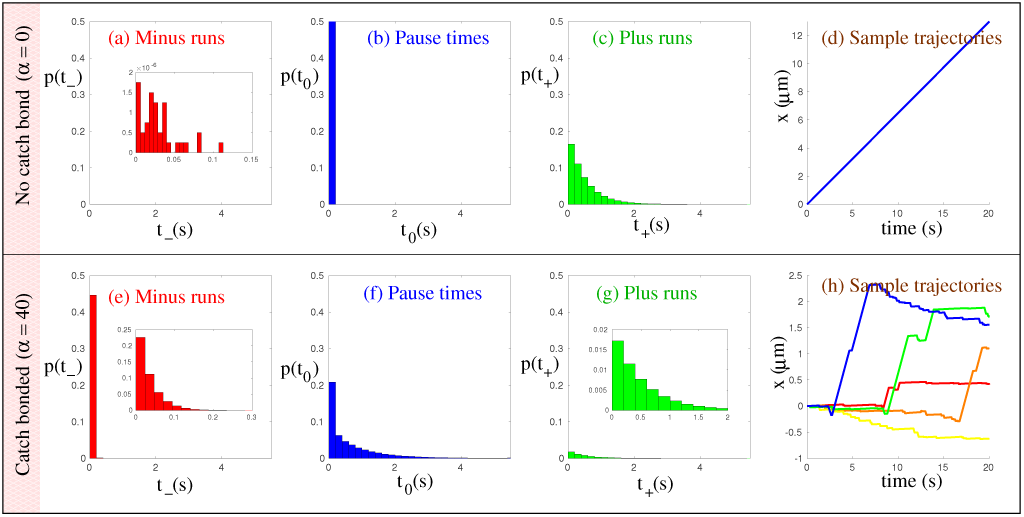
Probability distributions of runtimes for *N*_+_ = 6, *N*_−_ = 2. The top panels show the normalized histograms and sample trajectories for dynein in the absence of catch bond (*α* = 0). The bottom panels show the corresponding quantities for catch-bonded dynein (*α* = 40). (a) and (e) Distributions of runtimes for minus directed runs (shown in red); (b) and (f) pausetime distributions (shown in blue); (c) and (g) distributions of runtimes for minus directed runs (shown in green); and (d) and (h) sample trajectories. Insets, where present, show a magnified view of the probability distributions.

In *dictyostelium* cell extracts, it has been reported that teams of four to eight dyneins and one to two kinesins are simultaneously attached to a cargo^15^. The resultant motion was observed to be minus-end directed with intermittent pauses. To understand these results in the context of our model, we fixed *N*_+_ = 2 and *N*_−_ = 6 (Fig. 4). In the absence of catchbonding, the resultant motion is strongly plus-end directed (Fig. 4d). The probability distributions of runtimes show that there are many more kinesin runs (Fig. 4c) than dynein runs (Fig. 4a), and the average runtime is also higher in the case of kinesins. The pauses in this case are also of extremely short duration (Fig. 4b). Due to the stall force of kinesin motor being about 5 times that of dynein and π_0+_ > π_0−_ (see Table I), even with *N*_+_ < *N*_−_, a plus-end directed run, on average, continues for a longer time than a minus-end directed run, leading to larger average runtimes along the positive direction. When dynein catch bond is switched on, the picture changes dramatically. Minus-ended runs become much more frequent than plus-ended runs, while the average pause time also increases by an order of magnitude compared to the non-catchbonded case, and becomes comparable to the average minus directed runtimes. This is shown in Figs. 4(e)-(g). Load force on dynein due to attached kinesin engages the catch-bond, making it more difficult to unbind from the filament. Therefore, we see that the manifestation of catchbonding in dynein results in strong minus directed runs with longer duration pauses, as is shown in Fig. 4(h). This qualitatively agrees with the experimental observation of transport of endosomes in *Dictyostelium* cells^15^.

In a separate set of experiments on early endosomes in fungi, a team many kinesin motors (3-10) are in-volved in *tug-of-war* with 1 or 2 dynein motors during transport^5^. We generate the probability distribution of pause times along with minus-end and plus-end directed runtimes for a cargo being transported by 6 kinesins and 2 dyneins (*N*_+_ = 6, *N*_−_ = 2). The results displayed in Fig. 5 illustrates that while in the absence of catchbonding in dynein, the resultant motion would be strongly plus-end directed, with very small pause times, incorporation of catchbonding results in the frequency of minus-ended runs exceeding the frequency of plus-ended runs by almost one order of magnitude. However, the average duration of the minus-ended runs is about one order of magnitude lower than that of the plus-end directed run duration. Further there are now substantial duration of pauses (1–4 sec) during transport. These characteristics of the probability distributions result in typical cargo trajectories which exhibits bidirectional motion with pauses.

## DISCUSSION

In this article we have explicitly shown how incorporation of catchbonding behaviour of dynein motors in the modeling approach for bidirectional transport naturally leads to the phenomenon of codependent transport of cellular cargo. Many of the *in-vivo* and *in-vitro* experiments have characterized the nature of bidirectional transport in terms of cargo particle trajectories^5,15^. We qualitatively reproduce some of the experimentally observed codependent features seen in context of cargo trajectories during transport. The findings of our model points to the crucial role played by catchbonding in dynein motors in bidirectional transport, highlighting its significance as an internal regulatory mechanism during transport, albeit through mechanical interaction between the motors.

The effect of dynein motor inhibition on bidirectional transport was studied in *Drosophila* neurons through mutations in the dynein heavy chain (cDHC) and in the dynactin complex^17^. In the framework of our theory, this has been modeled through a decrease in the dynein number *N*_−_, or through varying *ε*_0−_. While both decreasing *N*_−_ or increasing *ε*_0−_ has the effect of weakening the dynein motor action, the manifestation of these two effects in the transport characteristics can in general be distinct. The results of these experiments can then qualitatively be understood in the light of Figs. 2(a) and 3(a), where weakening the dynein motor can lead to stalled motion of the cargo.

Diverse experiments have also indicated that mutations of conventional kinesin in Drosophila can hamper motion of cellular cargo in both directions^18,19,30–32^. This is consistent with the results shown in Figs. 2(b), where we show that reducing the number of kinesins can stall cargo motion completely. Interestingly, while kinesin exhibits a conventional slip bond, the cooperative force exerted by the catch bonded dynein on kinesins, and vice-versa, introduces a complex interplay which results in signatures of codependent transport being observed even on varying effective kinesin numbers. For example, as shown in Fig. 2(b) for weak dynein, reducing the number of kinesin, can in certain ranges, decrease the overall motion of the cargo in the negative direction. This counterintuitive phenomenon is a direct manifestation of the dynein catch bond. The introduction of a mechanical regulation mediated by the dynein catch bond, thus provides a plausible mechanism for codependent transport. Curiously enough, these processivity measures also point to the sharp difference in transport characteristics of strong dynein when compared to weak dynein. In the former case regulatory role of catchbonding is very weak since the typical force scale at which catchbond is activated is quite high with respect to the typical load forces experienced by the motors. It would indeed be interesting to probe further if this is the reason for the *strong* dynein in yeast not being involved in transport, while *weak* mammalian dynein are crucial to intracellular transport.

We also study the probability distribution of runtimes for plus-ended and minus-ended runs and the pause time distributions for our simulated cargo trajectories (Figs. 4 and 5), with and without catch bonds. We show that catchbonding dramatically alters the transport characteristics, and the nonlinear unbinding response of catch bonded dynein can provide one possible mechanism to understand qualitatively the observed experimental trajectories, in the context of two disparate experiments.

While our phenomenological model highlights the role of catchbonding as an internal regulatory mechanism for cellular transport, a complete description of bidirectional transport requires the incorporation of other experimental observations. First of all, some experiments have pointed to the presence of external regulatory mechanism which coordinates the action of the motors and the resultant transport of the cargo^1,3,19,33–39^. For instance, the protein JIP1 has been shown to regulate axonal transport^25^, while the *Klar* protein regulates transport of lipid droplets in *Drosophila*^1,3^. Another recent interesting observation has been that cargoes have *memory*, when plus moving motors detach they have a higher probability of moving in the opposite direction when they re-attach, and vice-versa^23^. Some recent experiments have also suggested that other factors such as the interactions between multiple motors leading to clustering, and the rotational diffusion of the cargo itself can play a role in regulating transport^40,41^. Further, our model makes a simplifying assumption in ignoring the stochastic load sharing by attached motors, as suggested by some experiments^22,42^. Finally, dynein motors exhibit variable step size under load^10,43^, which has not be taken into consideration in our theoretical description. However, even ignoring these various external regulatory factors, we show that codependent transport characteristics can arise simply from the incorporation of the dynein catch bond. A careful examination of the various experiments is required to delineate the relative importance of this internal regulatory mechanism that is mediated through catchbonding in motors and the various external modes of regulating transport.

In summary, our phenomenological model illustrates the key principle that the incorporation of a dynein catch bond should be an integral part of any theoretical description that attempts to explain bidirectional motion of cellular cargo. The model is able to capture the broad qualitative features observed in context of a multitude of experiments on motor driven transport within the cell. The framework proposed here encapsulates both the tug-of-war and codependent behaviour in appropriate regimes, and illustrates the importance of the dynein catch bond in determining the codependent transport characteristics.

## ACKNOWLEDGMENTS

MKM acknowledges financial support from the Ramanujan Fellowship (13DST052), DST India and the IRCC Seed Grant, IITB (14IRCCSG009). AC acknowledges SERB project No. EMR/2014/000791 for financial support. SM and MKM acknowledge SERB project No. EMR/2017/001335 for financial support. NG, SC and AC thank the HPC facility at IISER Mohali for computational time. SC and AC acknowledges DST, India for financial support. The authors thank the organizers of the SMYIM conference, Goa, and the ISPCM conference, Bangalore. MKM acknowledges the hospitality of MPIPKS, Dresden, where part of this work was done. SM and MKM acknowledge helpful discussions with Roop Mallik, TIFR.

## Appendix A: Materials and Methods

At any instant of time, the state of the cargo is characterised by the number of attached Kinesin (*n*_+_)and Dynein motors (*n*_−_). The maximum of number of kinesin and dynein motors are *N*_+_ and *N*_−_ respectively (0 < *n*_+_ < *N*_+_ and 0 < *n*_−_ < N_−_). The time evolution of the system is then governed by the master equation^9^

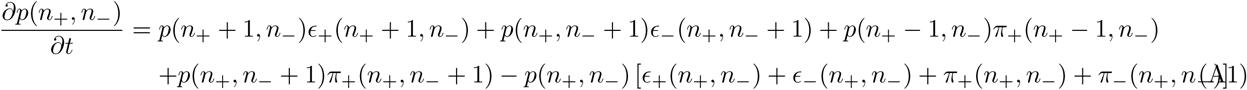

where, *p*(*n*_+_, *n*_–_) is the probability to find the cargo with *n*_+_ kinesin and *n*_−_ dynein motors.

The kinesin and dynein binding rates are assumed to be of the form π_±_ = (*N*_±_ – *n*_±_)π_0±_, where *N*_+_π_0+_ (*N*–π_0_–) is the rate for the first kinesin (dynein) motor to bind to the MT.

The unbinding rate for kinesin is given by the expression *ε*_+_ (*n*_+_,*n*_−_) = *n*_+_*ε*_0+_ exp[*F*_c_(*n*_+_,*n*_−_)/(*n*_+_*F*_d+_)], while the unbinding rate for dynein is given by *ε*_−_(*n*_+_,*n*_−_) = *n*_−_*ε*_0−_ exp[−*E_d_*(*F_c_*(*n*_+_,*n*_−_)) + *F_c_*(*n*_+_,*n*_−_)/(*n*_−_*F_d−_*)], with the catch bond deformation energy given by

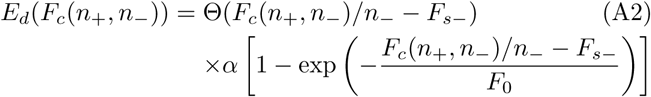

Here, *ε*_0±_ denotes the zero-force single motor unbinding rates, while *α* parameterizes the strength of the catch bond. The cooperative force felt by the motors due to the effect of the motors of the other species is given by^11^

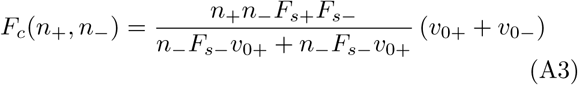

and the cargo velocity is given by

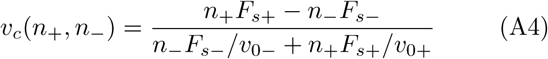

Here, υ_0±_ denotes the velocity of kinesin (or dynein) motors,

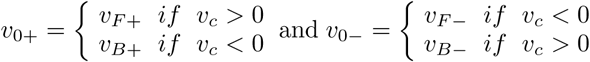

where, *υ_F_* and *υ_B_* are the forward and backward motor velocities. The stall forces for the two motor species are denoted by *F_s_*_±_.

The parameters used in the study are taken from the literature, and are summarized in Table I.

**TABLE I.** Single motor parameter values used in the simulations. The deformation force scale *F*_0_ is a phenomenological parameter, as determined in Ref^26^.

| Parameter | Kinesin | Ref. | Dynein | Ref. |
| --- | --- | --- | --- | --- |
| $F_{s\pm}$ | 6 pN | <sup>44</sup> | 1 pN (Weak)<br>7 pN (Strong) | <sup>45</sup><br><sup>46</sup> |
| $F_{d\pm}$ | 3 pN | <sup>44</sup> | 0.67 pN | <sup>10,26</sup> |
| $\pi_{0\pm}$ | 5/s | <sup>47</sup> | 1/s | <sup>48</sup> |
| $\epsilon_{0\pm}$ | 1/s | <sup>44</sup> | (0.1 - 10)/s | <sup>49</sup> |
| $F_0$ | NA | | 7 pN | <sup>10,26</sup> |
| $v_{F\pm}$ | 0.65 $\mu\text{m/s}$ | <sup>50</sup> | 0.65 $\mu\text{m/s}$ | <sup>51</sup> |
| $v_{B\pm}$ | 1 nm/s | <sup>50</sup> | 1 nm/s | <sup>52,53</sup> |

### 1. Numerical techniques

Time trajectories of cargo are obtained by simulating the master equation using the Stochastic Simulation Algorithm (SSA)^27,28^. All possible initial configurations were generated for a (*N*_+_, *N*_−_) pair, and 1000 trajectories were evolved for each initial configuration. A run finishes if the simulation continues until the maximum time *T_MAX_* or if all motors detach from the MT. The runlength was then averaged over all initial configurations and all iterations. Probability distributions were also computed from the SSA trajectories after discarding initial transients.

The probability distributions, and the motility diagrams, were also obtained by constructing the nullspace of the associated transfer matrix for the master equation. The probability distributions computed this method and the SSA algorithm matches exactly.

